# Variability of morphology-performance relationships under acute exposure to different temperatures in three strains of zebrafish

**DOI:** 10.1101/2023.12.18.572202

**Authors:** Christina L Miller, Robert J Dugand, Katrina McGuigan

## Abstract

Locomotion is thermally sensitive in ectotherms and therefore it is typically expressed differently among thermally heterogenous environments. Locomotion is a complex function, and while physiological and behavioural traits that influence locomotor performance may respond to thermal variation throughout life, other contributing traits, like body shape, may have more restricted responses. How morphology affects locomotor performance under variable temperature conditions is unknown. Here, we investigated three genetically distinct strains of zebrafish (AB, WIK, and Tu) with a shared multi-generational history at 28°C. After rearing fish at a constant 28°C, we measured prolonged swimming speed (*U*_crit_) at each of six temperatures (between 16°C and 34°C). Speed was strongly positively correlated among temperatures, resulting in most among-individual variation being temperature-independent (i.e., fish were relatively fast or slow across all temperatures). However, we also detected significant variation along two axes reflecting temperature-dependent variation, including one reflecting differences among individuals consistent with a hotter-colder tradeoff (i.e., individuals that were relatively fast at cooler temperatures were relatively slow at hotter temperatures and vice versa). Although strains differed in mean swimming performance, within strain (among-individual) patterns of speed variation were markedly consistent. Body shape and size explained significant variation among individuals in both temperature-independent and temperature-dependent axes of swimming speed variation. Notably, morphological traits that were most strongly associated with temperature-independent performance variation (i.e., faster-slower) differed from those associated with temperature-dependent (i.e., hotter-colder) variation. Further, there were significant differences among strains in both the direction and strength of association for specific morphological traits. Our results suggest that thermally heterogenous environments could have complex effects on the evolution of traits that contribute to whole organism performance traits.

## Introduction

Motile organisms rely on locomotion to undertake diverse activities, including finding food and mates. The repeated evolution of specific environment-locomotor performance relationships provides strong evidence for local adaptation of locomotor performance (e.g. Dalziel et al., 2012; Ghalambor et al., 2004; Hendry et al., 2011; Langerhans et al., 2004; McGuigan et al., 2003). An individual’s locomotor performance can also vary plastically in response to environmental factors. For example, speed typically varies with ambient temperature in ectotherms (e.g. Condon et al., 2010; Latimer et al., 2014; Logan et al., 2018). Temperature-dependent performance, characterised by thermal performance curves (TPCs: Huey & Kingsolver, 1989), is interpreted with respect to the thermal sensitivity of enzymatic reactions underpinning physiological traits (Angilletta et al., 2003; Hochachka et al., 2002). Temperature variation can occur over different spatial and temporal scales, including diurnal and seasonal changes, with physiological processes and locomotor performance similarly exhibiting rapid (<1hr) and more gradual (seasonal acclimation) responses to these changes in ambient temperature (Schulte et al., 2011; Seebacher et al., 2014).

Plastic responses to temperature change can depend on both the life-stage of the organism and the duration of exposure, as well as being trait dependent (Kellermann et al., 2019; Salinas et al., 2019). Morphology is typically sensitive to temperature during development (e.g. Elphick & Shine, 1998; Frazier et al., 2008; Sfakianakis et al., 2011), but, in contrast to physiological and behavioural traits, once developmental growth is completed, morphology typically does not respond to subsequent changes in temperature. Yet locomotor performance depends on morphological as well as physiological (and behavioural) traits. For example, fish body shape and size affects swimming speed via influences on drag and stability, while speed further depends on physiological traits, such as the contractile properties of muscle fibres (Videler, 1993). Morphological trait values and swimming speed are correlated among individuals (e.g. Conradsen & McGuigan, 2015; Hendry et al., 2011; Langerhans & Makowicz, 2009) and undergo correlated evolution under divergent selection regimes (e.g. Dalziel et al., 2012; Kern et al., 2016; McGuigan et al., 2003). If, and how, the contribution of physiology, morphology, and behaviour to locomotor performance changes with ambient temperature has received relatively little attention.

*Drosophila melanogaster* reared at colder temperatures develop larger wings and have better flight performance than warm-reared flies from the same population (Frazier et al., 2008), suggesting coordinated developmental plasticity of morphology and performance. In contrast, while both morphology and locomotor speed responded to rearing temperature in the lizard *Bassiana duperreyi*, the change in morphology did not explain the change in speed (Elphick & Shine, 1998). Notably, Kolok (1992) found that different traits explained variation in swimming speed among individual large-mouth bass (*Micropterus salmoides*) in summer versus winter, suggesting that the causal basis of inter-individual variation in locomotor performance might be seasonally dependent. Such heterogeneity in performance correlations might be important for adaptive evolution in natural populations. If selection acts on other traits through their contribution to locomotor performance (Arnold, 1983; Walker, 2007), then changes in the magnitude or sign of the correlation of those traits with locomotor speed will cause heterogeneity in selection on these causal traits, even when selection on the emergent function, locomotor speed, remains consistent.

Here, we investigate the relationship between morphology and swimming speed in zebrafish, *Danio rerio*. We specifically sought to determine whether body shape or size could explain the variation in the response of swimming speed to acute changes in temperature. Fish were reared to adulthood under constant thermal conditions of 28°C, and their swimming speed measured during acute (<4hr) exposure to six test temperatures: 16, 20, 24, 28, 31 and 34°C. We characterised the variation among individuals in their swimming speed across this temperature range, and determined whether morphology explained the observed inter-individual variation. We further determined how the relationships between static morphology and thermally dependent performance varied among three genetically differentiated populations (strains). Overall, our results suggest that morphology explains substantial variation in both temperature-independent and temperature-dependent swimming performance, and that these relationships are genetically variable.

## Methods

### Study system

Zebrafish, native to the Indian subcontinent, are a diurnal, shallow water schooling fish with a broad altitudinal and latitudinal distribution. Therefore, they occupy a wide thermal range (16°C to 34°C) and experience acute daily changes in temperature (Arunachalam et al., 2013; Engeszer et al., 2007; Spence et al., 2006). We assayed swimming speed and morphology of three widely studied strains: AB, WIK, and Tu. These strains have independent histories and are genetically differentiated (Holden & Brown, 2018; Suurväli et al., 2020), consistent with being collected from independent wild populations.

Local populations of the three strains were established by fish imported from the Zebrafish International Resource Center (ZIRC) three to five generations prior to this experiment. These stocks were maintained under very similar thermal conditions (28°C) as at ZIRC (28.5°C: Westerfield, 2007). To initiate the current experiment, clutches were collected from five breeding groups (each consisting of four males and four females) per strain, and reared in 3.5L tanks on a recirculating water system (for details on husbandry see Conradsen et al., 2016). Under these common-environment rearing conditions, phenotypic differences among strains can be attributed to genetic differences, while variation within strains will reflect both genetic and environmental (among and within rearing tank) effects, which cannot be partitioned under this breeding design.

From ∼90 days post fertilisation (dpf), sexually mature males were tagged using an elastomer tag (Northwest Marine Technology, Inc.) following protocols described in Conradsen and McGuigan (2015). These individually identifiable males were assigned to groups of six, with at least one fish per strain in each group, and were maintained with their group for the duration of the experiments. Males were assayed for their swimming performance and morphology across 150-264 dpf. As detailed below, we verified that males were close to their final (asymptotic) size at their first swimming trial.

### Swimming performance

Critical swimming speed, *U*_crit_, was assayed at 16°C, 20°C, 24°C, 28°C, 31°C, and 34°C using a step-velocity protocol (Brett, 1964) following protocols described in Conradsen and McGuigan (2015). Briefly, fish were swum in groups of six in a flume with a swim chamber of 46×14×13cm (LxWxH) (30L capacity, Loligo systems, Tjele, Denmark). Fish were allowed to settle at low flow (4cms^-1^) for 15 minutes. The flow rate was then increased by 4cms^-1^ every five minutes (300 seconds). Individual fish that could no longer hold station were removed, and their *U*_crit_ calculated as:

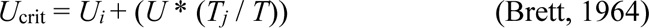

where *U_i_* was the maximum velocity (cm^-1^) maintained for the full step interval (*T*; here, 300 seconds), *U* was the water speed increment (here, 4cm^-1^), and *T_j_* was the time (in seconds) the individual fish swam at their final velocity step.

Water in the flume was maintained at temperatures 28°C and above using an aquarium tank heater (1500 W Titan Heavy Duty Aquarium Heater, Quian Hu, Singapore) and at temperatures 24°C and below using an aquarium tank chiller (440 W TECO Tank Chiller, Ravena Italy). Prior to swimming, fish were placed in tanks that were gradually cooled (at a rate of 0.20°C per minute) or warmed (0.27°C per minute) from 28°C to the test temperature. Our experimental design involved swimming each tank of six fish at each of the six temperatures, with groups encountering temperatures in a randomised order. Two flumes were initially used, but one malfunctioned, returning unreliable data. Data from this flume were discarded, and these assays were not repeated in the other flume, resulting in some loss of data (detailed below).

### Morphology

Fish were anesthetised (AQUI-S, Lower Hutt, New Zealand) and photographed on their left side using a tripod mounted Nikon Coolpix camera. A 1mm grid was included in all photographs. Images were randomly ordered, and the position of 12 landmarks recorded using TPSdig2 (Rohlf, 2005; Fig. 1). Fish size was recorded as standard length (SL), the distance from the anterior tip of the snout (LM1) to the origin of the caudal fin (LM6). Landmark positions were then aligned using generalised Procrustes fit, implemented in MorphoJ (Klingenberg, 2008; Klingenberg, 2011). Shape was characterised by 10 Inter-Landmark Distances (ILDs, Fig. 1) following Conradsen and McGuigan (2015) in units of centroid size, calculated as the Euclidean distance between aligned landmark co-ordinates using R (RStudio Core Team, 2019). Each fish was photographed three times: during the first (150 – 157 dpf), after the third (233 - 240 dpf) and after the final (257 - 264 dpf) set of swim trials. We analysed the average (over the three repeated measures) trait values per individual. On average, fish grew 0.5mm (where average SL was 28.1mm) between the first and second measures, and 0.04mm between the second and third. One individual was identified as a multivariate outlier based on Mahalanobis distance (*α =* 0.001, df = 11) (mahalanobis function in base R, RStudio Core Team, 2019) and excluded from morphological analyses.

**Fig. 1.**
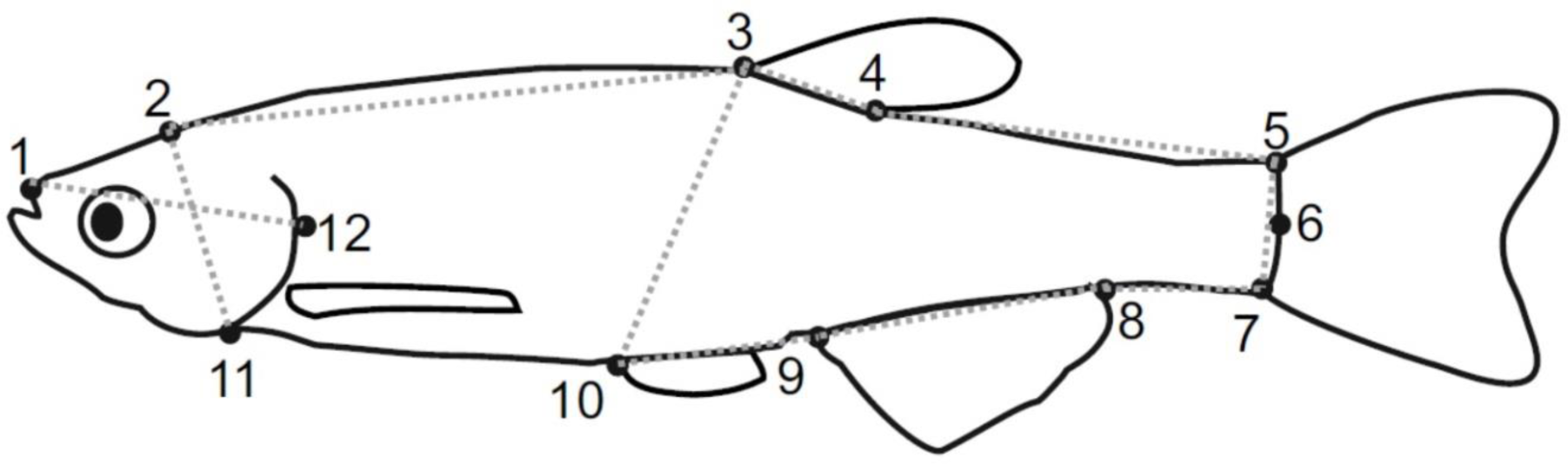
Schematic of *D. rerio* indicating the position of the 12 landmarks (black dots) and 10 inter-landmark distances (ILDS, grey dotted lines) used in this study. Landmarks were: anterior tip of the snout at the upper jaw (1); nape (2); dorsal-fin origin (3); dorsal-fin insertion (4); dorsal insertion of caudal fin (5); median caudal fin insertion (6); ventral insertion of caudal fin (7); ray of the anal fin (8); anal-fin origin (9), pelvic-fin origin (10); ventral posterior point of the operculum (11); posterior most point of the operculum (12). The 10 ILDs are referred to by their landmark endpoints (e.g., ILD1-12 is the distance between landmarks 1 and 12).

### Data analyses

#### Divergence among strains in thermal performance and morphology

*U*_crit_ was assayed for a total of 148 individuals (47, 48, and 53 of AB, Tu and WIK respectively), with 118 individuals measured across at least four temperatures. Data were missing at random with respect to temperature and strain, with a minimum (maximum) of 29 (40) individuals assayed per strain per temperature. Due to the missing data, we did not investigate multivariate outliers here. One (of 600) *U*_crit_ observation was extreme (> 3 SD above mean) and strongly inflated among-individual variance at that temperature; we therefore excluded this observation from all analyses.

We first aimed to determine whether the molecular genetic divergence among strains (Holden & Brown, 2018; Suurväli et al., 2020) was reflected in divergent means for speed and body shape. Although these strains are widely used, adult phenotypes have been compared only rarely, providing limited evidence of inter-strain divergence in comparable traits (Wakamatsu et al., 2019). With limited sample size (*n* = 3 strains) relative to the number of traits measured, sampling error could generate strong covariance of performance with morphology, and we therefore do not interpret strain-level morphology: performance relationships. Rather, we aim to assess these morphology: performance relationships among individuals within each strain, and to determine whether those relationships are divergent among strains.

There are several approaches for analysing thermal performance data. Our data were best suited to a multivariate approach (Gomulkiewicz et al., 2018; Latimer et al., 2014) because: individuals were assayed at a fixed set of temperatures; there were relatively few observations per individual (≤ 6); and there was only a small decline in performance across the hottest temperatures for some individuals. We determined whether strains differed in either their average response of speed to temperature (i.e., thermal performance) or in mean swimming speed. Maximum likelihood, implemented in PROC MIXED in SAS (SAS Institute Inc, 2011), was used to fit the model:

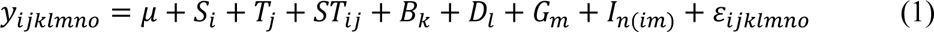

where *y* was the vector of *U*_crit_ measures (*o* = 1…599), and *μ* was the overall mean *U*_crit_. The effect of the *i^th^* (*i* = 1…3) Strain (*S*) and *j^th^*(*j* = 1…6) Temperature (*T*), along with their interaction (*ST_ij_*) were fit as categorical fixed effects. The effect of the *k*^th^ (*k* =1…6) swimming Block (*B*) and the *l^th^* (*l* = 1,2) Diurnal period (*D*, morning or afternoon) were also modelled as categorical fixed effects. Block (where fish were randomly allocated to a different temperature in each of the *k* = 1…6 repeated swimming blocks) captured variation due to age and experience, as well as potential temporal trends in general conditions. The effect of the *m^th^* (*m =* 1…29) swim Group (*G*) was modelled as a random effect to account for variation among mixed-strain groups of fish that were housed together and swam together. The effect of the *n*^th^ (*n* = 1…148) Individual (*I*), nested within strain, was also fit as a random effect, where the covariance among temperatures (repeated measures of the same individual at multiple temperatures), was constrained to be positive definite (all eigenvalues positive). The residual (ε) captured unexplained variation among the repeated measures per individual.

We applied log-likelihood ratio tests (LRT), comparing the fit (−2 log-likelihood) when the term of interest (the strain-by-temperature interaction or strain) was modelled versus when it was not; this difference in fit follows a chi-square distribution with the degrees of freedom equal to the number of parameters differing between the two models (here, 1) (Liang & Self, 1996). The LRT is robust to errors in identifying the appropriate degrees of freedom, given the complex error structure (repeated measures where sample sizes are unequal); conclusions were identical between this approach and a F-ratio test. Conclusions were also identical when assumptions about the rank of among-individual variance, homogeneity of among-individual variance among strains, and homogeneity of the residual were relaxed.

Next, we assessed whether strains differed in multivariate morphology. PROC GLM in SAS (SAS Institute Inc, 2011) was used to fit the model:

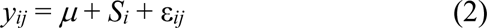

where *y* was a vector of the 11 morphological traits per individual, *μ* was the vector of global means per trait, and strain was modelled as a categorical fixed effect. Individual, nested within strain was the random residual error, ε. We retained the normalised linear discriminant functions from this analysis and calculated individual scores along each of the two discriminant functions for plotting. Inclusion of size as a response variable allowed us to succinctly determine whether strains were morphologically divergent, considering both size-dependent (allometric) and independent shape changes. However, these traits differ in scale (SL in cm versus ILDs in units of centroid size) which could influence conclusions. We applied model (2) to data on the raw measurement scale, and to variance standardised data (mean = 0, standard deviation = 1 for each trait). Conclusions (strains were morphologically distinct, with WIK and Tu differing the most) were consistent across both analyses, and we present only results on the variance standardised scale.

#### Performance: Morphology Relationships

We investigated the relationship between swimming performance and morphology with two aims: to determine whether morphology was associated with temperature-dependent and temperature-independent variation in *U*_crit_, and to determine whether these relationships differed among the strains.

We first defined temperature-dependent and temperature-independent swimming performance traits using the eigenvectors of the 6×6 matrix of among-individual variation, which has the variation in *U*_crit_ at each temperature along the diagonal, and covariance among temperatures off the diagonal. These eigenvectors reflect different patterns within the thermal performance variation (e.g., Izem & Kingsolver, 2005; Latimer et al., 2014). The covariance matrix was estimated using restricted maximum likelihood (REML) to fit a modified form of model (1) containing only random effects. Data were centred (mean = 0) on their respective level of each of the categorical fixed effects prior to analysis, which is equivalent to fitting these in the model, but improves efficiency; these mean-centred data were retained to calculate performance scores (detailed below). The among-individual variance was sequentially modelled (using a type = FA0(*n*) statement) as having variance (i.e. eigenvalues > 0) in *n* = 0 up to *n* = 6 independent traits (i.e. eigenvectors); LRT were applied to compare these nested models to determine how many axes of variation were statistically supported (Hine & Blows, 2006; Meyer & Kirkpatrick, 2005), and thus how many eigenvectors we would retain as performance traits. Consistent with individual speed being moderately correlated across temperatures, three independent performance axes (i.e., the eigenvectors; Table 1) were statistically supported, and these defined our thermal performance trait set. We detail the interpretation of these as temperature-dependent or independent in the Results. Note that there was little evidence that strains differed in this among-individual variation (Supplementary Materials; Table S1, S2), and we therefore utilise all data to estimated the common among-individual variance.

**Table 1.**
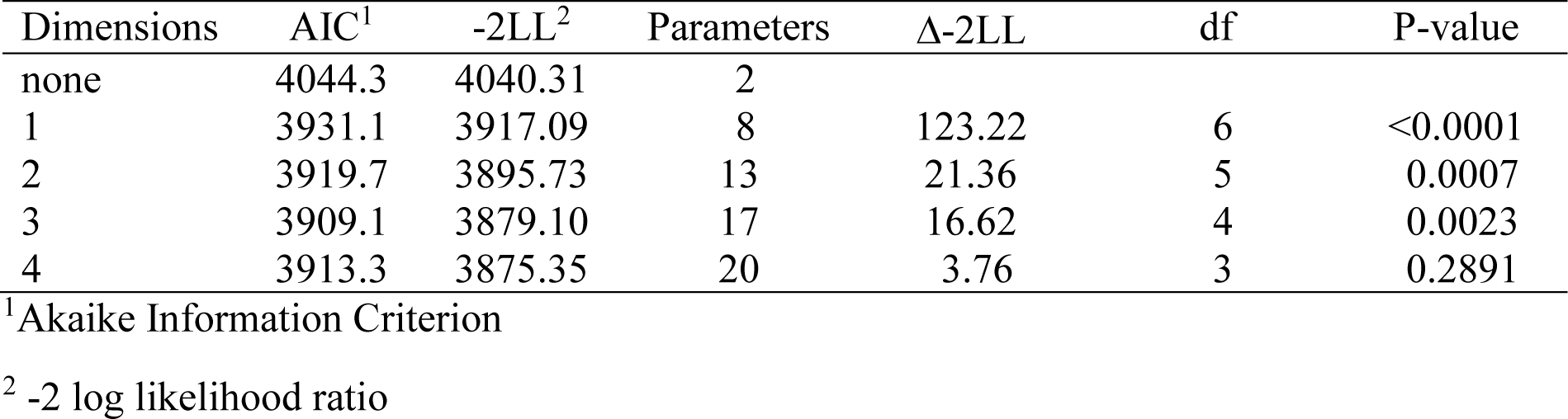
Dimensionality of among-individual thermal performance. The fit (AIC; −2LL), number of estimated parameters, and results of the likelihood ratio tests are reported for models with none through four dimensions (i.e., eigenvalues > zero) of among-individual variation in swimming speed across the six temperatures. The difference in fit (Δ-2LL) between a model and the next dimension model (none to 1; 1 to 2, etc.) follows a chi-square distribution with degrees of freedom equal to the difference in number of parameters.

| Dimensions | AIC<sup>1</sup> | -2LL<sup>2</sup> | Parameters | Δ-2LL | df | P-value |
| --- | --- | --- | --- | --- | --- | --- |
| none | 4044.3 | 4040.31 | 2 |  |  |  |
| 1 | 3931.1 | 3917.09 | 8 | 123.22 | 6 | <0.0001 |
| 2 | 3919.7 | 3895.73 | 13 | 21.36 | 5 | 0.0007 |
| 3 | 3909.1 | 3879.10 | 17 | 16.62 | 4 | 0.0023 |
| 4 | 3913.3 | 3875.35 | 20 | 3.76 | 3 | 0.2891 |

<sup>1</sup>Akaike Information Criterion
<sup>2</sup> -2 log likelihood ratio

To calculate individual scores on the three performance axes (using the mean-centred data), we first excluded individuals with fewer than four *U*_crit_ observations, retaining 118 individuals (AB = 40, Tu = 40, WIK = 38). Missing data were replaced with the mean swimming speed deviation of that strain at the missing temperature(s). These data were then multiplied by their respective loadings on the eigenvectors of among-individual variation in *U*_crit_. Because performance scores were calculated on mean-centred data, the *R*^2^ values from regression of performance on morphology are expected to be upwardly biased, akin to the known inflation of heritability estimates in mixed models including fixed effects (Wilson, 2008). Notably, *a priori* accounting for some sources of variation in speed data could not generate spurious associations between speed and the morphological variables, which were independently collected and processed.

For each independent performance axis, we first tested the hypothesis that the three strains shared the same morphology-performance relationship. Strains differed (see Results) in both their thermal performance curves (and hence scores on the performance axes), and in average morphology. To consider the relationship between morphology and performance separately from these differences, and to report standardised regression coefficients for comparison, we first centred and scaled (i.e., calculated z-scores) the data within each strain. We fit two linear regression models (using the lm function in R RStudio Core Team, 2019): one with the 11 morphological traits as continuous predictors, and a second in which strain-by-morphological trait interactions were also fit. These models were compared via LRT (implemented in the lmtest package, Zeileis & Hothorn, 2002). Where the null hypothesis of among-strain homogeneity of slopes was rejected, the emmeans package (Lenth, 2023) was used to further investigate differences.

Given evidence of strain-specific slopes (see Results), we analysed data within each strain to determine the best model to explain thermal performance variation. Using the ‘regsubsets’ function in the ‘leaps’ package in R (RStudio Core Team, 2019), an exhaustive search was made of all possible additive linear models, including from one up to all 11 mophological trait predictors. The 10 best-fit (based on residual sums of squares) models were retained for each model size (i.e., number of predictors, with only one 11-trait model; 101 models were retained per performance trait and strain). We identified the best-fit model based on Mallow’s Cp criterion. To support robust interpretation of differences among strains, we also report the model averaged regression coefficients from a subset of models within 2 AIC (Aikike Information Criterion) of this best-fit model (implemented using the R package ‘MuMIn’ Bartoń, 2022).

## Results

### Divergence among strains

Strains differed in how their mean swimming speed changed with temperature (i.e., their thermal performance: strain-by-temperature interaction, *X*^2^ = 29.98, df = 1, *p* < 0.0001; Fig. 2). WIK had the slowest speed at all temperatures, although the magnitude of the difference was not constant (Fig. 2). Tu was markedly faster than either WIK or AB at the coldest temperatures, while AB was fastest at the hottest temperatures (although AB and Tu had similar speeds at these temperatures) (Fig. 2).

**Fig. 2.**
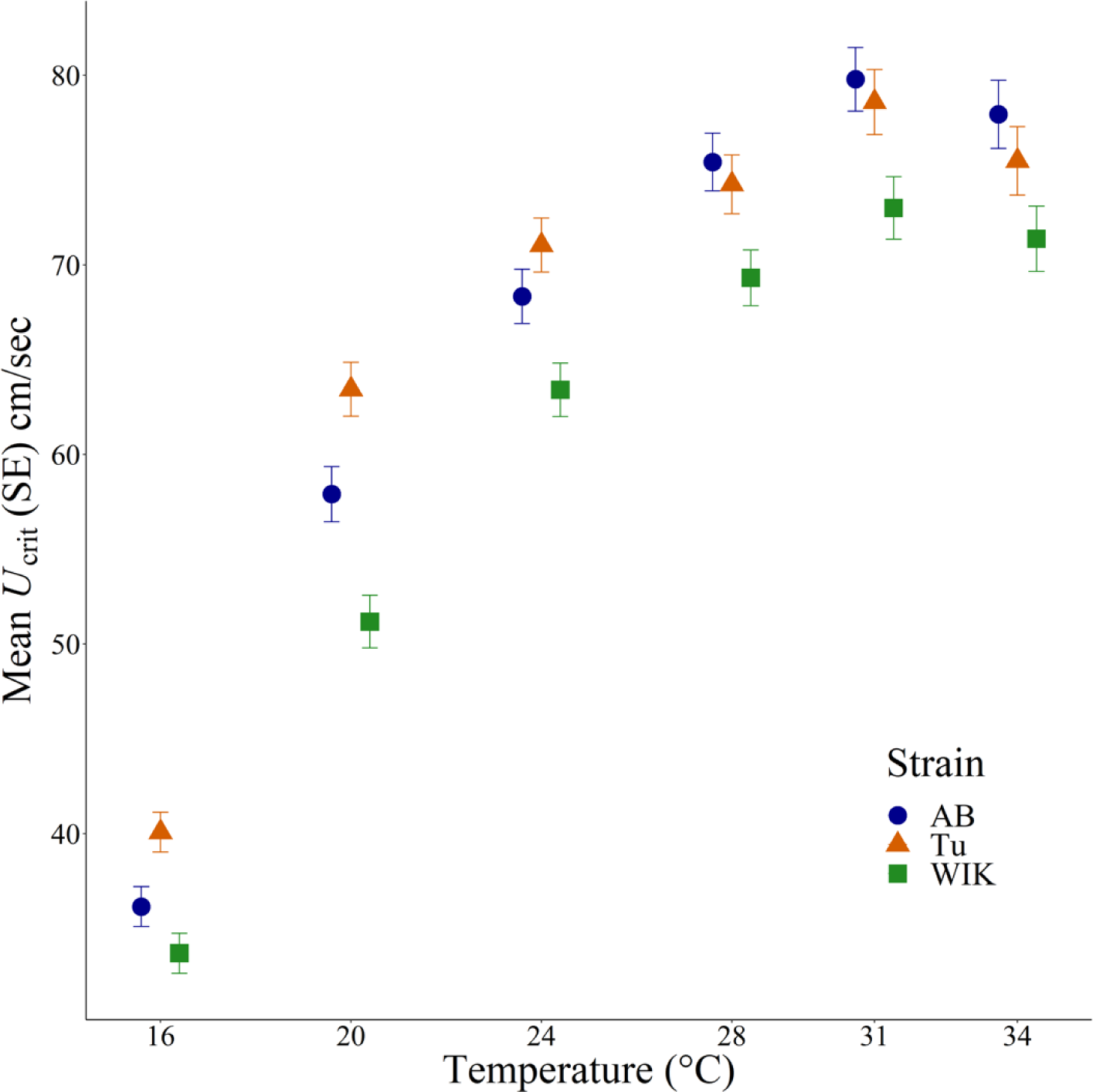
Swimming speed variation of three zebrafish strains across six temperatures. The least square means (± SE) from model (1) are plotted for each strain. The points are offset to allow a visualisation of each strain at each temperature.

Strains also differed significantly in their morphology (Wilk’s λ = 0.09, *F*_22, 260_ = 28.01, *p* < 0.0001). WIK and Tu were the most strongly differentiated, with the higher mean *DFS*_1_ score of Tu (Fig. 3) reflecting their longer (SL), shallower bodies (ILD 3.10) (*DFS*_1_: Table 2). AB was differentiated from the other strains along the second discriminant axis (Fig. 3), reflecting a relatively short dorsal caudal peduncle (ILD 4.5) (*DFS*_2_: Table 2).

**Fig. 3.**
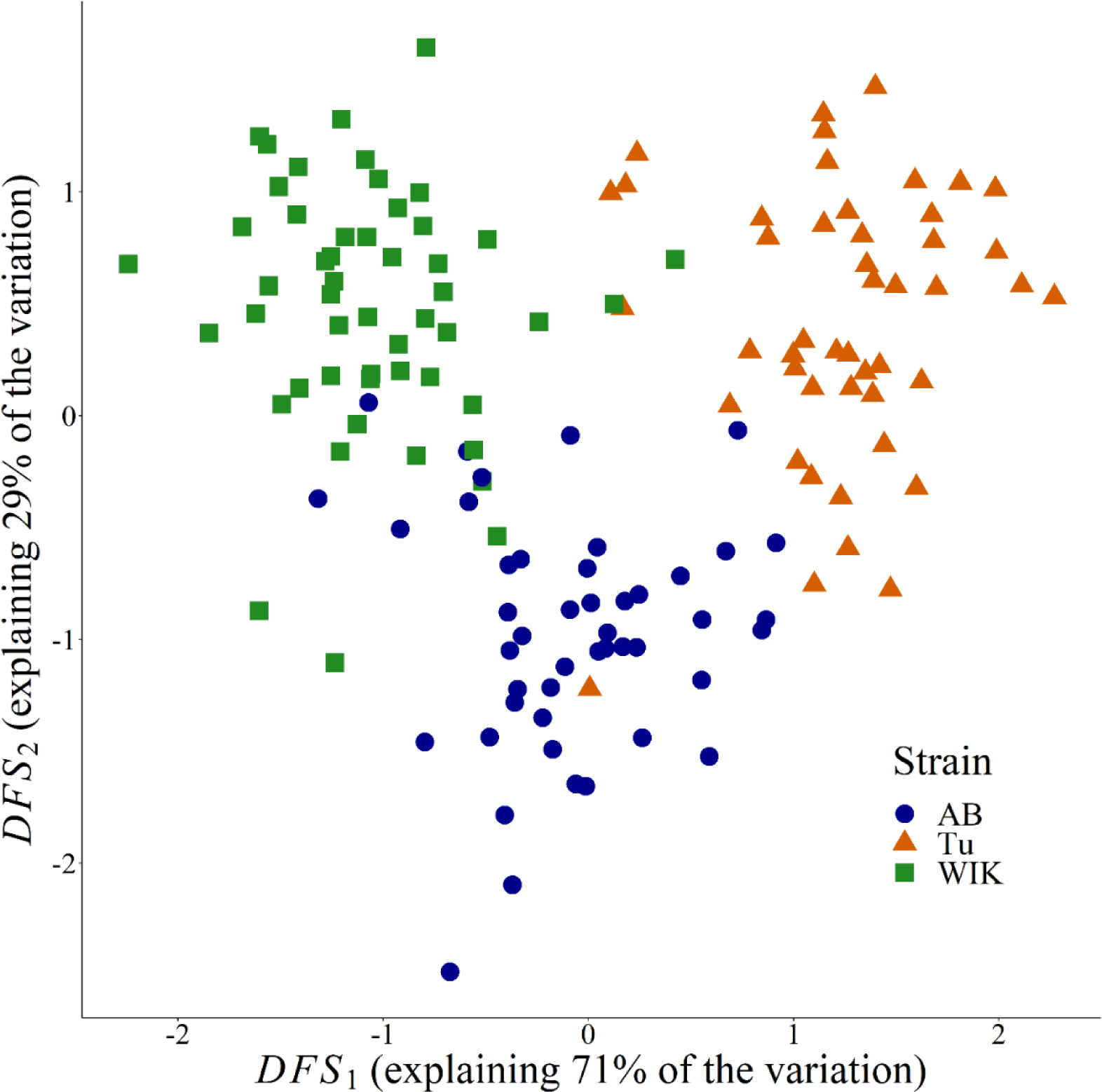
Variation in morphology among strains. Individual scores along the two discriminant functions are plotted by strain. The percent of among-strain variation is shown on the respective axis.

**Table 2.**
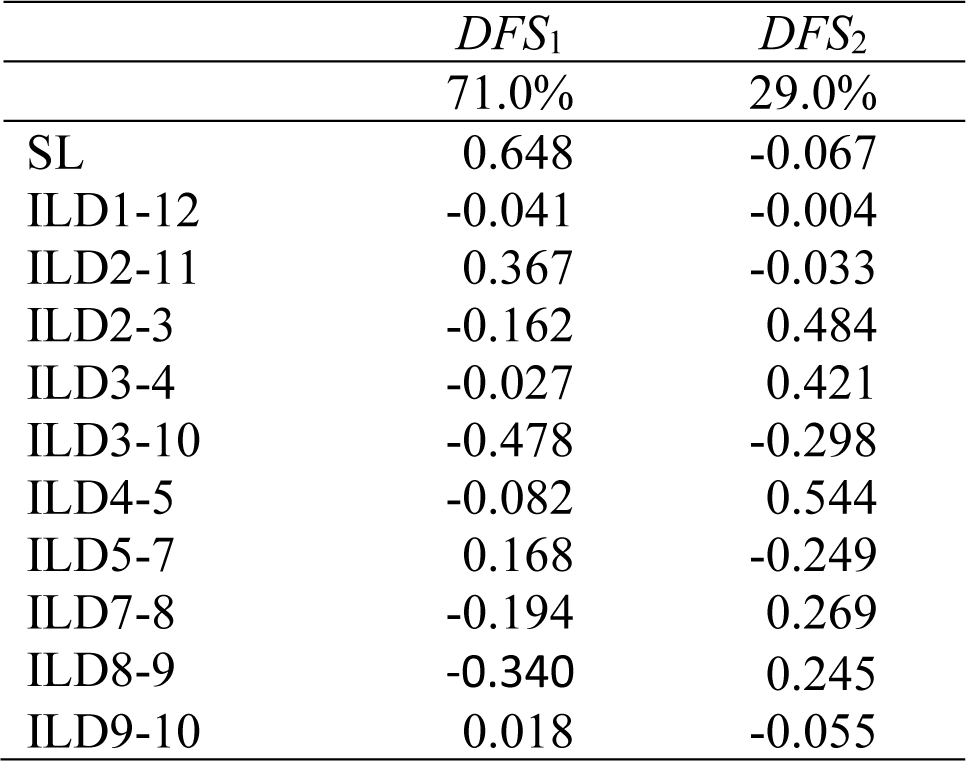
Discriminant functions of morphological variation among strains. The proportion of among-strain variance (%) accounted for, and normalised trait loadings are presented for the two among-strain linear discriminant functions (*DFS*_1_ and *DFS*_2_). Traits are defined in Fig. 1.

### Variation among individuals in swimming performance

All temperatures contributed to the major axis of among-individual variation in swimming performance in the same direction, with relatively similar magnitude of loadings (with the weakest and strongest contribution from 16°C and 31°C, respectively, reflecting heterogeneity in the magnitude of among-individual variances at each temperature: diagonals of Table S1) (*e*_1_: Table 3). This pattern reflects the pervasive positive cross-temperature correlations of speed (off-diagonal elements of Table S1) and suggests that most among-individual variation in speed was independent of temperature (i.e., a faster-slower axis of variation).

**Table 3.** Among-individual variation in swimming performance. The covariance matrix of among-individual variation in *U*_crit_ at the six assay temperatures (for all strains pooled, constrained to three dimensions; Table S1) was subject to eigenanalysis. The eigenvalues (top row; as a proportion of total variation in second row) and normalised eigenvector loadings of for each temperature are reported.

| | $e_1$ | $e_2$ | $e_3$ |
| --- | --- | --- | --- |
|  | 135.63 | 31.08 | 19.37 |
|  | 72.9% | 16.7% | 10.4% |
| 16°C | 0.138 | -0.267 | -0.021 |
| 20°C | 0.439 | -0.436 | -0.087 |
| 24°C | 0.392 | -0.463 | -0.074 |
| 28°C | 0.431 | -0.111 | -0.103 |
| 31°C | 0.563 | 0.689 | -0.384 |
| 34°C | 0.363 | 0.193 | 0.910 |

The second axis of among-individual variation in performance was characterised by contrasting contributions from speed at cool (20°C and 24°C) and warm (31°C) temperatures (*e*_2_: Table 3), somewhat consistent with a hotter-colder pattern of variation: individuals that were relatively fast at colder temperatures were relatively slow at hot temperatures (particularly 31°C) (and vice versa). The third axis was dominated by speed at 34°C (*e*_3_: Table 3), which contributed little to the other two axis of variation. Speed at 34°C was relatively weakly correlated with speed at other temperatures (Table S1), and thus *e*_3_ captures the independent determination of speed at this, hottest, temperature.

### Morphology-performance relationships

For *e*_1_, which predominantly reflected temperature-independent (faster-slower) variation in speed, morphology explained significant variation in all three strains (Table 4), but the morphology-performance relationships were strain-specific (LRT of model with versus without strain-specific slopes: *X*^2^ = 42.69, df = 22, *p* = 0.0052; Table S3). Caudal peduncle depth (ILD5-7) had the greatest difference in effect, strongly positively associated with higher *e*_1_ scores (i.e., faster swimming) in Tu, but with a weak negative effect on speed in AB and no effect in WIK (Table 5, Table S4). Speed increased with greater distance between pelvic and anal fin origins (ILD9-10) in both Tu and AB, but this trait again did not influence *e*_1_ scores in WIK (Table 4, Table S4). Longer heads (ILD1-12) were strongly associated with slower swimming in Tu, but weakly implicated as increasing speed in the other strains (Table 5, Table S4). In WIK, speed increased with standard length (SL), but was independent of length in the other two strains (Table 5, Table S4).

**Table 4.** Regression of swimming performance on morphology. Results for the regression model with the lowest Mallow’s *Cp* are reported for each strain and performance axis (regression coefficients for included morphological variables are reported in Table 5).

| | $e_1$ | $F$ | df | $p$ | Adj $R^2$ |
| --- | --- | --- | --- | --- | --- |
| **AB** |  | 3.62 | 4,35 | 0.0144 | 0.212 |
| **Tu** |  | 11.81 | 4,35 | <0.0001 | 0.526 |
| **WIK** |  | 10.41 | 2,34 | 0.0003 | 0.343 |
| | $e_2$ | | | | |
| **AB** |  | 3.89 | 3,36 | 0.0166 | 0.182 |
| **Tu** |  | 5.99 | 3,36 | 0.0020 | 0.277 |
| **WIK** |  | 3.29 | 3,33 | 0.0169 | 0.196 |
| | $e_3$ | | | | |
| **AB** |  | 1.19 | 1,38 | 0.1748 | 0.023 |
| **Tu** |  | 4.98 | 3,36 | 0.0055 | 0.234 |
| **WIK** |  | 1.01 | 1,35 | 0.3239 | <0.001 |

**Table 5.** Regression coefficients from the best fit model of swimming performance on morphology. For each axis of performance variation (*e*_1_, *e*_2_, *e*_3_), the standardised beta coefficients, β (± SE), are reported for strain-specific (*e*_1_, *e*_2_) or experiment wide (*e*_3_) models. Morphological traits are defined in Fig. 1; performance traits are defined in Table 3. Where the null hypothesis of zero slope was rejected (α = 0.05) values are shown in bold.

| | $e_1$ | | | $e_2$ | | | $e_3$ |
| --- | --- | --- | --- | --- | --- | --- | --- |
|  | AB | Tu | WIK | AB | Tu | WIK | Tu |
| SL |  |  | **0.551** (0.135) |  |  |  |  |
| ILD1-12 |  | **-0.566** (0.148) |  |  |  |  | -0.286 (0.166) |
| ILD2-11 |  | -0.297 (0.156) |  | **0.368** (0.149) |  | 0.316 (0.179) |  |
| ILD2-3 |  |  |  |  | **-0.330** (0.147) |  |  |
| ILD3-4 |  |  |  | **-0.400** (0.154) |  |  | **0.501** (0.184) |
| ILD3-10 | 0.423 (0.271) |  |  |  |  |  |  |
| ILD4-5 | **0.566** (0.212) |  | 0.242 (0.135) |  |  | -0.290 (0.165) | **0.486** (0.201) |
| ILD5-7 | -0.334 (0.168) | **0.512** (0.121) |  |  |  | **-0.428** (0.168) |  |
| ILD8-9 |  |  |  | 0.240 (0.150) | -0.273 (0.144) |  |  |
| ILD9-10 | 0.346 (0.186) | **0.286** (0.116) |  |  | **0.385** (0.139) |  |  |

There was also evidence that morphology explained significant variation along the second performance axis, *e*_2_, in each strain (Table 4), but that strains differed in how morphology influenced this performance (LRT: *X*^2^ = 37.25, df = 22, *p* = 0.0222). This overall support for strain-specific morphology-performance relationships reflected relatively weak differences in slope of individual traits (Table S3). Specifically, Tu fish with high *e*_2_ scores (i.e., fish that swam relatively fast at 31°C but relatively slowly at 20°C and 24°C) had relatively anteriorly positioned dorsal fins (i.e., short ILD2-3), but this trait had a weakly opposing effect on *e*_2_ performance in WIK and no effect in AB (Table 5, Table S5). In WIK, fish with shallow caudal peduncles (ILD5-7) had high *e*_2_ scores, while in AB there was a weakly positive slope, and no effect in Tu (Table 5, Table S5).

For *e*_3_ (predominantly reflecting variation in speed at 34°C), there was no evidence that strains differed in how morphology influenced scores (*X*^2^ = 21.50, df = 22, *p* = 0.4902), but morphology did influence *e*_3_ scores (pooled analysis of all strains: *F*_1,115_ = 4.14, *p* = 0.0443, adjusted *R*^2^ = 0.026). Only one trait was included in the best-fit model – higher *e*_3_ scores (i.e., faster speed at 34°C) were associated with shorter heads (ILD1-12: *β* = −0.186, SE = 0.092). These results suggests that, although individuals varied significantly along this performance axis (Table S2), very little of that variation could be explained by morphological variation. Notably, while there was no statistical support for heterogeneity of slopes between strains, morphology explained significant variation along *e*_3_ in only Tu (Table 4). Tu fish with higher *e*_3_ scores had shorter heads (consistent with the pooled model) and longer posterior dorsal lengths (ILD3-4 and ILD4-5: Table 5).

## Discussion

Locomotor performance is complex, influenced by morphological, behavioural, and physiological traits that are themselves determined by genes and environment. Temperature is well known exert immediate effects on locomotor performance via effects on the contractile properties of skeletal muscle (Angilletta et al., 2002; James & Tallis, 2019). Temperature can also elicit plastic changes in behaviour, which may either compensate for or exacerbate the potential fitness consequences of temperature effects on performance (reviewed in James & Tallis, 2019). How morphology, which does not respond to acute temperature changes, contributes to variation in performance under thermally variable conditions remains relatively unexplored. Examining variation in body shape and in prolonged swimming speed across a temperature gradient within three genetically distinct strains of zebrafish we found that external morphology explained patterns of both temperature-independent and temperature-dependent performance. The morphological trait associations were specific both to the axis of thermal performance variation, and to the genotype (strain).

Given the broad literature reporting correlated evolution (e.g. Cano-Barbacil et al., 2020; Kern et al., 2016; Langerhans, 2009b) or within population covariation (e.g. Conradsen et al., 2016; Hendry et al., 2011) of morphology and speed measured at a single temperature, we expected body shape to predict temperature-independent speed. Consistent with this, morphology explained variation among-individuals on the major axis of performance variation, *e*_1_, an axis that most closely resembled a faster-slower (Izem & Kingsolver, 2005) mode of thermal performance. In contrast, we expected that thermally dependent performance variation would be due to variation in musculature and enzymes that were not reflected in the external morphological traits under analysis. However, body shape also explained significant variation among individuals along *e*_2_, a performance axis which reflected a hotter-colder (Izem & Kingsolver, 2005) pattern where individuals that were relatively fast at cooler temperatures (20°C and 24°C) tended to be relatively slow at hotter temperatures (31°C) and vice versa.

Within each strain, the different axes of swimming performance variation were typically associated with different morphological traits, with little evidence that the same trait had either concordant or antagonistic effects on temperature-independent (*e*_1_) and temperature-dependent (*e*_2_) axes of variation. Indeed, when considering the three strains and two major axes of swimming performance, all 11 morphological traits were represented in at least one best-fit model, yet only ventral inter-fin length (ILD9-10) in Tu was associated with higher performance scores on both axes, while in WIK dorsal caudal length (ILD4-5) had weakly antagonistic effects on the two performance axes (Table 5). Below, we discuss three broad factors that are important for interpreting these results.

Firstly, we note that the mechanisms that may connect static morphology to dynamic performance are not clearly established. The viscosity of water decreases with increasing temperature, which may affect swimming independently of the physiological effects of temperature (Danos & Lauder, 2012; Fuiman & Batty, 1997; von Herbing & Keating, 2003). While viscosity has the greatest impact on small (larval) fishes (Fuiman & Batty, 1997; von Herbing & Keating, 2003; Yavno & Holzman, 2018), complex effects on larger fishes (including adult zebrafish) have been documented (Danos & Lauder, 2012). Temperature-dependent physiological rates and viscosity can both influence tail beat frequency and amplitude (Danos & Lauder, 2012; Fuiman & Batty, 1997), which may change recoil dynamics during swimming. Notably, although strains differed in their morphology-*e*_2_ associations (Table 5, Table S5), in general, implicated traits capture changes in positioning of dorsal and paired fins, which may influence recoil (yaw) (Borazjani, 2013; Conradsen & McGuigan, 2015; Webb, 2006). Numerical simulations have extended our understanding of the mechanisms of swimming (Gazzola et al., 2014; Tokić & Yue, 2012). Simulations exploring fine-scale (intra-specific) variation, coupled with further empirical experiments, could provide novel insights into the contribution of external morphology to locomotor performance, and how this might vary with environmental effects.

Secondly, we emphasise that while regression tests causal hypotheses (i.e., body shape causes performance values) it does not provide evidence of causation. Manipulative experiments can be effective at confirming causative effects – for example, experimental fin clips have supported the causal role of fins in determining swimming speed (e.g. Wakamatsu et al., 2019). However, such experiments are impossible or impractical for most traits. Replicated natural co-evolution of trait and performance (e.g. Langerhans, 2008) are suggestive of a causal effect, but may arise from responses to independent selection pressures. Correlated evolution of performance in response to artificial selection on a putatively causal trait (e.g. Kern et al., 2016) may provide more compelling inference. Such an approach has not yet been applied to the question of how body shape influences thermally dependent swimming speed.

Finally, morphological covariance with independent axes of thermal performance variation could influence interpretations of adaptive evolution. Many studies have demonstrated broadly consistent patterns of phenotypic evolution among locomotor niches (e.g. Langerhans, 2008, 2009), suggesting a common morphology-performance relationship. However, these studies also typically reveal unique responses in different populations inhabiting broadly similar environments (Heckley et al., 2022; Langerhans, 2018). Including thermal variation in habitat descriptions, alongside well-studied factors such as water flow, structural complexity, and predation, may improve our understanding of locomotor performance evolution.

Studies of traits influencing locomotor performance predominantly consider taxa (populations or species) with divergent trait values. To predict future evolutionary responses, we need to better understand the factors determining within population variation. Our observation of consistent ranking of fast versus slow individuals, irrespective of temperature, (i.e., along *e*_1_) suggests that performance capacity is repeatable; repeatability sets an upper limit on the evolutionary potential of a trait (Boake, 1989; Wilson, 2018). Previous studies, at a single temperature, have demonstrated that locomotor performance is moderately repeatable over various timescales (reviewed in Conradsen et al., 2016). However, repeatability (or heritability) of temperature-dependent locomotor performance has received less attention (Latimer et al., 2014; Logan et al., 2018). Our observation that *e*_2_, reflecting temperature-dependent (hotter-colder) variation in *U*_crit_, was associated with similar variation in the three, genetically divergent, strains (Table S2) suggests that it also captures repeatable differences among individuals. In contrast, the statistical support for strain-specific morphological predictors of performance suggests that the morphology-performance map may be more variable than performance itself.

Notably, strains differed in the contribution of caudal peduncle depth (ILD5-7) to among-individual differences in speed. The caudal region plays a key role in determining swimming performance, with hydrodynamic principals (Blake, 2004; Walker, 1997) and empirical studies (e.g. Langerhans, 2008; Langerhans et al., 2004) revealing that larger caudal regions generate greater thrust but increased drag, increasing the maximum speed, but decreasing endurance. Conradsen et al. (2016) observed strong, positive slopes of caudal peduncle depth on *U*_crit_ at 28°C in repeated measures (∼3 months apart) of both male and female WIK zebrafish. Here, WIK caudal peduncle depth was not associated with overall speed (*e*_1_ scores), although fish with relatively deep caudal peduncles had low *e*_2_ scores (i.e., fish that were relatively fast at 20°C and 24°C but slow at 31°C had deeper caudal peduncles: Table 5). The morphology-performance map of Tu for *e*_1_ (Tables S4) closely matched that of the most similar-age WIK fish in the previous study (normalised vector dot product = 0.79, where 1.0 would indicate concordance and 0.0 orthogonal vectors) (Table S4 versus Assay 1 in Table 2 of Conradsen et al., 2016). Thus, our results provided equivocal evidence that morphological traits had repeatable associations with the repeatable axes of locomotor performance.

In conclusion, our study has provided some evidence that static morphology may contribute to variation in thermally dependent swimming speed. We suggest that both modelling approaches and incorporation of data on thermal regime into studies of natural divergence in locomotor performance may improve our understanding of causal effects on locomotor capacity. We also highlight the paucity of data on within-population variation in locomotion and its putatively causative traits, and the consequent limited knowledge of what proportion of observed phenotypic variation reflects heritable genetic differences among individuals. Ongoing statistical developments and increasing accessibility of genomic data may provide opportunities to investigate the genetic covariance of performance and morphological traits in natural populations.

## Supporting information

Supplementary Methods and Results

## References

Angilletta, M. J., Niewiarowski, P. H., & Navas, C. A. (2002). The evolution of thermal physiology in ectotherms. Journal of Thermal Biology, 27, 249–268. 10.1016/S0306-4565(01)00094-8

Angilletta, M. J., Wilson, R. S., Navas, C. A., & James, R. S. (2003). Tradeoffs and the evolution of thermal reaction norms. Trends in Ecology & Evolution, 18(5), 234–240. 10.1016/s0169-5347(03)00087-9

Arnold, S. J. (1983). Morphology, performance, and fitness.. American Zoologist, 23(2), 347–361.

Arunachalam, M., Raja, M., Vijayakumar, C., Malaimmal, P., & Mayden, R. (2013). Natural history of zebrafish (*Danio rerio*) in India. Zebrafish, 10(1), 1–14. 10.1089/zeb.2012.0803

Bartoń, K. (2022). MuMIn: Multi-Model Inference. R package version 1.47.1. https://CRAN.R-project.org/package=MuMIn

Blake, R. W. (2004). Fish functional design and swimming performance. Journal of Fish Biology, 65, 1193–1222. 10.1111/j.1095-8649.2004.00568.x, availableonlineat http://www.blackwell-synergy.com

Boake, C. R. B. (1989). Repeatability: Its role in evolutionary studies of mating behaviour. Evolutionary Ecology, 3, 173–182.

Borazjani, I. (2013). The functional role of caudal and anal/dorsal fins during the C-start of a bluegill sunfish. Journal of Experimental Biology, 216(9), 1658–1669. 10.1242/jeb.079434

Brett, J. R. (1964). The respiratory metabolism and swimming performance of young sockeye salmon. Journal of the Fisheries Research Board of Canada, 21(5), 1183–1226. 10.1139/f64-103

Cano-Barbacil, C., Radinger, J., Argudo, M., Rubio-Gracia, F., Vila-Gispert, A., & Garcia-Berthou, E. (2020). Key factors explaining critical swimming speed in freshwater fish: a review and statistical analysis for Iberian species. Scientific Reports, 10(1), 18947. 10.1038/s41598-020-75974-x

Condon, C. H., Chenoweth, S. F., & Wilson, R. S. (2010). Zebrafish take their cue from temperature but not photoperiod for the seasonal plasticity of thermal performance. Journal of Experimental Biology, 213(21), 3705–3709. 10.1242/jeb.046979

Conradsen, C., & McGuigan, K. (2015). Sexually dimorphic morphology and swimming performance relationships in wild-type zebrafish *Danio rerio*. Journal of Fish Biology, 87(5), 1219–1233. 10.1111/jfb.12784

Conradsen, C., Walker, J. A., Perna, C., & McGuigan, K. (2016). Repeatability of locomotor performance and morphology-locomotor performance relationships. Journal of Experimental Biology, 219(18), 2888–2897. 10.1242/jeb.141259

Dalziel, A. C., Vines, T. H., & Schulte, P. M. (2012). Reductions in prolonged swimming capacity following freshwater colonization in multiple threespine stickleback populations. Evolution, 66(4), 1226–1239. 10.1111/j.1558-5646.2011.01498.x

Danos, N., & Lauder, G. V. (2012). Challenging zebrafish escape responses by increasing water viscosity. Journal of Experimental Biology, 1(215), 1854–1862. 10.1242/jeb.068957

Elphick, J. M., & Shine, R. (1998). Longterm effects of incubation temperatures on the morphology and locomotor perfromance of hatchling lizards (*Bassiana duperreyi*, Scincidae). Biological Journal of the Linnean Society, 63, 429–447. 10.1111/j.1095-8312.1998.tb01527.x

Engeszer, R. E., Patterson, L. B., Rao, A. A., & Parichy, D. M. (2007). Zebrafish in the wild: a review of natural history and new notes from the field. Zebrafish, 4(1), 21–40. 10.1089/zeb.2006.9997

Frazier, M. R., Harrison, J. F., Kirkton, S. D., & Roberts, S. P. (2008). Cold rearing improves cold-flight performance in *Drosophila* via changes in wing morphology. Journal of Experimental Biology, 211(13), 2116–2122. 10.1242/jeb.019422

Fuiman, L., & Batty, R. (1997). What a drag it is getting cold: partitioning the physical and physiological effects of temperature on fish swimming. Journal of Experimental Biology, 200(12), 1745–1755. 10.1242/jeb.200.12.1745

Gazzola, M., Argentina, M., & Mahadevan, L. (2014). Scaling macroscopic aquatic locomotion. Nature Physics, 10, 758–761. 10.1038/NPHYS3078

Ghalambor, C. K., Reznick, D. N., & Walker, J. A. (2004). Constrains on adaptive evolution: the functional trade-off between reproduction and fast-start swimming performance in the Trinidadian Guppy (*Poecilia reticulata*). The American Naturalist, 164, 38–50. 10.1086/421412

Gomulkiewicz, R., Kingsolver, J. G., Carter, A., & Heckman, N. (2018). Variation and evolution of function-valued traits.. *Annual Review of Ecology*, Evolution and Systematics, 49, 139–164. doi10.1146/annurev-ecolsys-110316-022830

Heckley, A. M., Pearcem A. E., Gotanda, K. M., Hendry, A. P., & Oke, K. B. (2022). Compiling forty years of guppy research to investigate the factors contributing to (non)parallel evolution. Journal of Evolutionary Biology, 35(11), 1414–1431. 10.1111/jeb.14086

Hendry, A. P., Hudson, K., Walker, J. A., Rasanen, K., & Chapman, L. J. (2011). Genetic divergence in morphology-performance mapping between Misty Lake and inlet stickleback. Journal of Evolutionary Biology, 24(1), 23–35. 10.1111/j.1420-9101.2010.02155.x

Hine, E., & Blows, M. W. (2006). Determining the effective dimensionality of the genetic variance-covariance matrix. Genetics, 173(2), 1135–1144. 10.1534/genetics.105.054627

Hochachka, P. W., Hochachka, P. W., & Somero, G. N. (2002). Biochemical adaptation: Mechanism and process in physiological evolution. Oxford University Press.

Holden, L. A., & Brown, K. H. (2018). Baseline mRNA expression differs widely between common laboratory strains of zebrafish. Scientific Reports, 8(1), 4780. 10.1038/s41598-018-23129-4

Huey, R. B., & Kingsolver, J. G. (1989). Evolution of thermal sensitivity of ectotherm performance. Trends in Ecology & Evolution, 4(5), 131–135.

Izem, R., & Kingsolver, J. G. (2005). Variation in continuous reaction norms: Quantifying directions of biological interest. American Naturalist, 166(2), 277–289. 10.1086/431314

James, R. S., & Tallis, J. (2019). The likely effects of thermal climate change on vertebrate skeletal muscle mechanics with possible consequences for animal movement and behaviour. Conservation Physiology, 7(1), coz066. 10.1093/conphys/coz066

Kellermann, V., Chown, S. L., Schou, M. F., Aitkenhead, I., Janion-Scheepers, C., Clemson, A., Scott, M. T., & Sgrò, C. M. (2019). Comparing thermal performance curves across traits: how consistent are they? Journal of Experimental Biology, 222(11). 10.1242/jeb.193433

Kern, E. M. A., Robinson, D., Gass, E., Godwin, J., & Langerhans, R. B. (2016). Correlated evolution of personality, morphology and performance. Animal Behaviour, 117, 79–86. 10.1016/j.anbehav.2016.04.007

Klingenberg, C. P. (2008). MorphoJ. Faculty of Life Sciences, University of Manchester.

Klingenberg, C. P. (2011). MorphoJ: An integrated softwear package for geometric morphometrics. Molecular Ecology Resources, 11, 353–357. 10.1111/j.1755-0998.2010.02924.x

Kolok, A. S. (1992). Morphological and physiological correlates with swimming performance in juvenile largemouth bass. American Journal of Physiology, 263(32), 1042–1048. 10.1152/ajpregu.1992.263.5.R1042

Langerhans, R. B. (2008). Predictability of phenotypic differentiation across flow regimes in fishes. Integrative and Comparative Biology, 48(6), 750–768. 10.1093/icb/icn092

Langerhans, R. B. (2009). Morphology, performance, fitness: functional insight into a post-Pleistocene radiation of mosquitofish. Biology Letters, 5(4), 488–491. 10.1098/rsbl.2009.0179

Langerhans, R. B. (2009b). Trade-off between steady and unsteady swimming underlies predator-driven divergence in *Gambusia affinis*. Journal of Evolutionary Biology, 22(5), 1057–1075. 10.1111/j.1420-9101.2009.01716.x

Langerhans, R. B. (2018). Predictability and parallelism of multitrait adaptation. Journal of Heredity, 109(1), 59–70. 10.1093/jhered/esx043

Langerhans, R. B., Layman, C. A., Shokrollahi, A. M., & DeWitt, T. J. (2004). Predator-driven phenotypic diversification in *Ganbusia affins*. Evolution, 58(10), 2305–2318. 10.1111/j.0014-3820.2004.tb01605.x

Langerhans, R. B., & Makowicz, A. M. (2009). Shared and unique features of morphological differentiation between predator regimes in Gambusia caymanensis. Journal of Evolutionary Biology, 22(11), 2231–2242. 10.1111/j.1420-9101.2009.01839.x

Latimer, C. A., McGuigan, K., Wilson, R. S., Blows, M. W., & Chenoweth, S. F. (2014). The contribution of spontaneous mutations to thermal sensitivity curve variation in *Drosophila serrata*. Evolution, 68(6), 1824–1837. 10.1111/evo.12392

Lenth, R. V. (2023). emmeans: Estimated Marginal means, aka Least-Squares means. R package version 1.8.4-1. https://CRAN.R-project.org/package=emmeans

Liang, K.-Y., & Self, S. G. (1996). On the asymptotic behaviour of the pseudolikelihood ratio test statistic. Journal of the Royal Statistical Society, 58(4), 785–796. 10.1111/j.2517-6161.1996.tb02116.x

Logan, M. L., Curlis, J. D., Gilbert, A. L., Miles, D. B., Chung, A. K., McGlothlin, J. W., & Cox, R. M. (2018). Thermal physiology and thermoregulatory behaviour exhibit low heritability despite genetic divergence between lizard populations. Proceedings of the Royal Society B, 285(1878), 1–14. 10.1098/rspb.2018.0697

McGuigan, K., Franklin, C. E., Moritz, C., & Blows, M. W. (2003). Adaptation of rainbow fish to lake and stream habitats. Evolution, 57(1), 104–118. 10.1111/j.0014-3820.2003.tb00219.x

Meyer, K., & Kirkpatrick, M. (2005). Restricted maximum likelihood estimation of genetic principal components and smoothed covariance matrices. Genetics Selection Evolution, 37(1), 1–30. 10.1186/1297-9686-37-1-1

Rohlf, F. J. (2005). *tpsDig2*. Department of Ecology and Evolution, State University of New York.

RStudio Core Team. (2019). R: A language and environment for statistical computing. In R Foundation for Statistical Computing. https://www.R-project.org/

Salinas, S., Irvine, S. E., Schertzing, C. L., Golden, S. Q., & Munch, S. B. (2019). Trait variation in extreme thermal environments under constant and fluctuating temperatures. Philosophical Transactions Of The Royal Society, 374, 20180177. 10.1098/rstb.2018.0177

SAS Institute Inc. (2011). The SAS system for Windows. In (Version version 9.4)

Schulte, P. M., Healy, T. M., & Fangue, N. A. (2011). Thermal performance curves, phenotypic plasticity, and the time scales of temperature exposure. Integrative and Comparative Biology, 51(5), 691–702. 10.1093/icb/icr097

Seebacher, F., White, C. R., & Franklin, C. E. (2014). Physiological plasticity increases resilience of ectothermic animals to climate change. Nature Climate Change, 5(1), 61–66. 10.1038/nclimate2457

Sfakianakis, D. G., Leris, I., Laggis, A., & Kentouri, M. (2011). The effect of rearing temperature on body shape and meristic characters in zebrafish (Danio rerio) juveniles. Environmental Biology of Fishes, 92(2), 197–205. 10.1007/s10641-011-9833-z

Spence, R., Fatema, M. K., Reichard, M., Huq, M. A., Wahab, M. A., Ahmed, Z. F., & Smith, C. (2006). The distribution and habitat preferences of the zebrafish in Bangladesh. Journal of Fish Biology, 69(3), 1435–1448. 10.1111/j.1095-8649.2006.01206.x

Suurväli, J., Whiteley, A. R., Zheng, Y., Gharbi, K., Leptin, M., & Wiehe, T. (2020). The laboratory domestication of zebrafish: From diverse populations to inbred substrains. Molecular Biology Evolution, 37(4), 1056–1069. 10.1093/molbev/msz289

Tokić, G., & Yue, D. K. P. (2012). Optimal shape and motion of undulatory swimming organisms. Proceedings of the Royal Society B, 279(1749), 3065–3074. 10.1098/rspb.2012.0057

Videler, J. J. (1993). Fish Swimming. Chapman and Hall.

von Herbing, I., & Keating, K. (2003). Temperature-induced changes in viscosity and its effects on swimming speed in larval haddock. The Big Fish Bang: Proceedings of the 26th Annual Larval Fish Conference.

Wakamatsu, Y., Ogino, K., & Hirata, H. (2019). Swimming capability of zebrafish is governed by water temperature, caudal fin length and genetic background. Scientific Reports, 9(1), 16307. 10.1038/s41598-019-52592-w

Walker, J. A. (1997). Ecological morphology of lacustrine threespine stickleback *Gasterosteus aculeatus* L. (Gasterosteidae) body shape. Biological Journal of the Linnean Society, 61, 3–50. 10.1111/j.1095-8312.1997.tb01777.x

Walker, J. A. (2007). A general model of functional constraints on phenotypic evolution. American Naturalist, 170(5), 681–689. 10.1086/521957

Webb, P. W. (2006). Stability and maneuverability. Elsevier Academic Press.

Westerfield, M. (2007). The zebrafish book: A guide for the laboratory use of zebrafish (Danio rerio). University of Oregon Press.

Wilson, A. J. (2008). Why *h^2^* does not always equal *V_A_/V_P_*? Journal of Evolutionary Biology, 21, 647–650. 10.1111/j.1420-9101.2008.01500.x

Wilson, A. J. (2018). How should we interpret estimates of individual repeatability? Evolution Letters, 2(1), 4–8. 10.1002/evl3.40

Yavno, S., & Holzman, R. (2018). Do viscous forces affect survival of marine fish larvae? Revisiting the ‘safe harbour’ hypothesis. Reviews in Fish Biology and Fisheries, 28, 201–212. 10.1007/s11160-017-9503-0

Zeileis, A., & Hothorn, T. (2002). Diagnostic checking in regression relationships. R News, 2(3), 7–10. https://CRAN.R-project.org/doc/Rnews/

