## Supplementary Methods and Results for "Variability of morphology-performance relationships under acute exposure to different temperatures in three strains of zebrafish"

To ensure that our approach to defining axes of swimming performance for further analysis did not generate differences among strains, we explicitly assessed whether strains differed in their among-individual variation in $U_{crit}$. The model fit to centred data to determine dimensionality and major axes of among-individual variance in swimming performance was constrained to estimate homogeneous variance across all three strains (main text Methods). For the statistically supported three dimensional [TYPE = FA0(3)] model, we compared the fit (-2LL) of this constrained model to a model fitting strain-specific among-individual covariance (implemented using the GROUP statement). There was weak statistical evidence that the strains differed in their among-individual variation (LRT of constrained and strain-specific models: $X^2 = 43.30$, df = 30, $P = 0.0551$). The among-individual covariance matrices for the strains pooled, and each individual strain, are shown in Table S1.

We further investigated how among-individual swimming performance variation differed among strains using Krzanowski's common subspace analysis (Aguirre et al., 2014; Krzanowski, 1979). The Krzanowski analysis considered the three statistically supported eigenvectors of the among-individual covariance matrices estimated for each strain. The eigenvalues of the Krzanowski subspace matrix (**H**) are bounded between 0 and $p$, where 0 indicates no shared orientation and $p$ is the number of matrices compared; here $p = 3$ strains. An eigenvalue of 3 indicates that the associated eigenvector of **H** is represented in the covariance matrix of each strain, while eigenvalues < 3 indicate at least one strain does not have a completely coincident axis of variation. To determine which strains differed, and how, we projected the normalised (unit length) eigenvectors of **H**, and of the pooled and strain-specific covariance matrices, through each estimated matrix (Aguirre et al., 2014).

The first eigenvalue of the Krzanowski's common subspace, 2.96, very close to the maximum of 3.0. This reflected the sharing among all three strains of the same 1st eigenvector of the within-strain covariance matrix (Table S2). The other two eigenvalues (2.82 and 2.43) of the common subspace were lower, but still broadly consistent with at least two strains sharing a common orientation of variation. Relative to its own 2nd eigenvector, AB had markedly reduced variance along the 2nd eigenvector of among-individual variation of the other strains, particularly WIK (Table S2). AB also had considerably lower total among-individual variance (trace: 155, 243 and 244 for AB, Tu and WIK, respectively; Table S1). Thus, the weak evidence of divergence among strains ($P$ =0.0551 as presented above) was primarily due to divergence of AB from Tu and WIK in both magnitude and orientation of variation.

As expected, given the weak statistical evidence of strain divergence, differences among strains were relatively small, the weight of evidence suggests there was substantial among-individual variation that was common (i.e., that there was non-zero among-individual variance in each strain along each eigenvector of the common subspace: Table S2). We therfore focus our interpretation of how temperature affected $U_{\text{crit}}$ (and its relationship with morphology) on the pooled estimate, which takes advantage of the total data.

### References


Aguirre, J. D., Hine, E., McGuigan, K., & Blows, M. W. (2014). Comparing G: multivariate analysis of genetic variation in multiple populations. *Heredity*, *112*(1), 21-29. https://doi.org/10.1038/hdy.2013.12

Houle, D., & Meyer, K. (2015). Estimating sampling error of evolutionary statistics based on genetic covariance matrices using maximum likelihood. *J Evol Biol*, *28*(8), 1542-1549. https://doi.org/10.1111/jeb.12674

Krzanowski, W. J. (1979). Between-group comparison of principal components. *Journal of the American Statistical Association*, *74*(367), 703-707.

RStudio Core Team. (2019). *R: A language and environment for statistical computing*. In R Foundation for Statistical Computing. https://www.R-project.org/

Venables, W. N., & Ripley, B. D. (2002). *Modern applied statistics with S.* (Fourth Edition ed.).

**Table S1. Among individual covariance in $U_{crit}$ among temperatures.** The among-individual variance in swimming speed at each temperature is on the diagonal, with covariances and correlations between temperatures below and above the diagonal, respectively. The REML-MVN CI are shown (in italics) below each estimate. Analyses were conducted on data for all strains pooled, or with among-individual covariances estimated separately for each strain (AB, Tu and WIK). The total variance (sum of the size variances, i.e., the matrix trace; CI in brackets) is given for each dataset. Each of the estimates are from models constrained to three dimensions (i.e., three positive eigenvalues). We took a REML-MVN approach (Houle & Meyer, 2015) to place confidence intervals on parameter estimates. We used the MASS package (Venables & Ripley, 2002) in R (RStudio Core Team, 2019) to generate 10,000 samples from the multivariate normal (MVN) distribution for pooled data (All Strains) and for each strain, using the REML covariance estimates to define the mean, and the REML inverse of the Fisher information matrix to define the variance. Because the REML covariance parameter estimates were constrained to have positive eigenvalues, the REML-MVN CIs may be upwardly biased; all hypothesis testing was therefore based on LRTs.

| | | | | | | |
|---|---|---|---|---|---|---|
| | | | **All Strains** Trace: 186.08 (*183.72, 868.84*) | | | |
| | 16°C | 20°C | 24°C | 28°C | 31°C | 34°C |
| 16°C | **4.79** | 0.95 | 0.97 | 0.81 | 0.29 | 0.37 |
| | ***1.73, 9.41*** | *0.70, 1.00* | *0.69, 1.00* | *0.30, 0.97* | *-0.17, 0.61* | *-0.09, 0.60* |
| 20°C | 11.85 | **32.21** | 1.00 | 0.95 | 0.56 | 0.52 |
| | *5.70, 19.04* | ***23.10, 48.49*** | *0.87, 1.00* | *0.50, 0.99* | *0.13, 0.69* | *-0.07, 0.70* |
| 24°C | 11.19 | 29.76 | **27.64** | 0.92 | 0.50 | 0.49 |
| | *5.42, 17.90* | *20.99, 44.14* | ***20.16, 42.59*** | *0.47, 0.98* | *0.13, 0.66* | *-0.25, 0.64* |
| 28°C | 9.00 | 27.35 | 24.68 | **25.78** | 0.79 | 0.62 |
| | *3.60, 15.64* | *18.19, 41.65* | *16.65, 38.67* | ***21.22, 73.89*** | *0.51, 0.95* | *0.17, 0.83* |
| 31°C | 4.94 | 24.82 | 20.57 | 31.27 | **60.56** | 0.54 |
| | *-4.98, 15.06* | *11.31, 43.29* | *9.33, 41.17* | *21.36, 201.52* | ***57.23, 466.36*** | *0.28, 0.71* |
| 34°C | 4.82 | 17.49 | 15.25 | 18.77 | 25.09 | **35.11** |
| | *-2.40, 13.66* | *-5.47, 43.66* | *-21.86, 29.25* | *8.27, 80.61* | *16.45, 176.56* | ***31.48, 262.90*** |
| | | | **AB** Trace: 155.15 (*145.36, 496.45*) | | | |
| | 16°C | 20°C | 24°C | 28°C | 31°C | 34°C |
| 16°C | **4.52** | 0.91 | 0.99 | 0.37 | 0.33 | 0.39 |
| | ***0.63, 11.72*** | *0.37, 1.00* | *0.63, 1.00* | *-0.30, 0.81* | *-0.29, 0.77* | *-0.12, 0.70* |

| | | | | | | |
|---|---|---|---|---|---|---|
| 20°C | 9.39 | **23.79** | 0.96 | 0.72 | 0.70 | 0.50 |
| | *1.70, 20.29* | ***12.63, 49.31*** | *0.71, 1.00* | *0.16, 0.91* | *0.20, 0.88* | *-0.32, 0.80* |
| 24°C | 12.68 | 28.16 | **36.33** | 0.50 | 0.47 | 0.44 |
| | *2.93, 24.73* | *15.15, 54.96* | ***22.59, 70.48*** | *-0.02, 0.78* | *0.03, 0.72* | *-0.33, 0.72* |
| 28°C | 3.29 | 14.89 | 12.65 | **17.92** | 0.99 | 0.57 |
| | *-3.91, 12.15* | *3.87, 34.07* | *-0.69, 31.49* | ***12.39, 82.75*** | *0.81, 1.00* | *0.09, 0.85* |
| 31°C | 4.52 | 21.81 | 17.95 | 26.86 | **40.71** | 0.48 |
| | *-5.89, 17.25* | *7.38, 49.06* | *1.51, 43.20* | *16.95, 145.76* | ***32.45, 191.61*** | *0.02, 0.75* |
| 34°C | 4.70 | 13.76 | 14.88 | 13.58 | 17.24 | **31.88** |
| | *-1.78, 14.35* | *-15.31, 39.81* | *-24.03, 38.38* | *2.87, 58.50* | *0.91, 69.75* | ***23.70, 142.90*** |

**Tu** Trace: 242.91 (*218.89, 470.37*)

| | 16°C | 20°C | 24°C | 28°C | 31°C | 34°C |
|---|---|---|---|---|---|---|
| 16°C | **3.29** | 0.96 | 0.68 | 0.46 | -0.10 | 0.16 |
| | ***0.27, 9.53*** | *0.41, 1.00* | *-0.08, 1.00* | *-0.34, 0.93* | *-0.75, 0.61* | *-0.40, 0.62* |
| 20°C | 11.70 | **45.40** | 0.87 | 0.69 | 0.19 | 0.26 |
| | *1.53, 24.78* | ***27.20, 96.47*** | *0.60, 0.99* | *0.29, 0.90* | *-0.21, 0.55* | *-0.22, 0.64* |
| 24°C | 7.81 | 36.69 | **39.56** | 0.93 | 0.66 | 0.38 |
| | *-0.63, 20.49* | *19.82, 80.57* | ***25.69, 80.71*** | *0.66, 0.99* | *0.28, 0.86* | *-0.09, 0.72* |
| 28°C | 5.33 | 29.30 | 37.01 | **40.16** | 0.80 | 0.65 |
| | *-5.15, 18.09* | *10.80, 70.32* | *20.99, 83.22* | ***27.93, 89.44*** | *0.41, 0.95* | *0.20, 0.92* |
| 31°C | -1.35 | 10.02 | 31.74 | 38.86 | **58.75** | 0.39 |
| | *-18.87, 12.54* | *-13.06, 40.30* | *13.25, 77.00* | *19.38, 96.91* | ***46.00, 147.37*** | *0.00, 0.71* |
| 34°C | 2.10 | 12.92 | 17.81 | 30.66 | 22.58 | **55.76** |
| | *-7.58, 12.07* | *-13.31, 49.95* | *-4.92, 53.48* | *9.79, 78.21* | *0.27, 70.39* | ***38.62, 115.18*** |

**WIK** Trace: 243.57 (*235.36, 1123.46*)

| | 16°C | 20°C | 24°C | 28°C | 31°C | 34°C |
|---|---|---|---|---|---|---|
| 16°C | **11.41** | 0.96 | 1.00 | 0.93 | 0.25 | 0.44 |

| | | | | | | |
|---|---|---|---|---|---|---|
| | ***3.42, 24.01*** | *0.64, 1.00* | *0.75, 1.00* | *0.37, 0.99* | *-0.25, 0.61* | *-0.11, 0.75* |
| 20°C | 19.95 | **38.19** | 0.98 | 0.83 | 0.49 | 0.67 |
| | *7.80, 35.31* | ***24.33, 63.72*** | *0.78, 1.00* | *0.30, 0.95* | *-0.02, 0.72* | *0.15, 0.83* |
| 24°C | 15.91 | 28.59 | **22.36** | 0.91 | 0.33 | 0.51 |
| | *6.09, 28.74* | *16.28, 49.48* | ***13.09, 40.84*** | *0.42, 0.98* | *-0.10, 0.61* | *-0.06, 0.75* |
| 28°C | 20.81 | 34.30 | 28.55 | **44.32** | 0.23 | 0.18 |
| | *8.12, 36.96* | *15.11, 66.21* | *14.63, 58.82* | ***33.75, 152.08*** | *-0.37, 0.55* | *-0.69, 0.46* |
| 31°C | 8.19 | 29.23 | 14.84 | 15.00 | **92.15** | 0.75 |
| | *-14.01, 30.29* | *-2.10, 74.73* | *-7.29, 54.93* | *-86.44, 53.14* | ***82.14, 697.94*** | *0.48, 0.91* |
| 34°C | 8.73 | 24.63 | 14.32 | 6.92 | 42.92 | **35.15** |
| | *-3.05, 24.22* | *8.68, 48.64* | *-2.51, 32.13* | *-132.12, 21.79* | *28.14, 341.73* | ***27.69, 211.28*** |

**Table S2. Among-individual variance associated with vectors of variation in thermal swimming performance.** Each of the eigenvectors of the Krzanowski’s common subspace matrix (**H**), and the first three eigenvectors of the pooled and strain specific among-individual covariance matrices were projected through the pooled and strain-specific covariance matrices (and each of their 10,000 REML-MVN samples, obtained as described in Table S1) to estimate the magnitude of variance in that direction of performance space. The variance (length) estimates of each vector in each covariance matrix are shown in bold, with the 90% CI (from the 10,000 REML-MVN matrices) below in italics. The length of the vectors through their own matrix (grey shaded entries) represent the eigenvalues of the those eigenvectors.

| | $h_1$ | $h_2$ | $h_3$ | Pooled 1 | Pooled 2 | Pooled 3 | AB1 | AB2 | AB3 | Tu1 | Tu2 | Tu3 | WIK1 | WIK2 | WIK3 |
|---|---|---|---|---|---|---|---|---|---|---|---|---|---|---|---|
| Pool | **135.01** | **21.89** | **28.06** | **135.63** | **31.08** | **19.37** | **133.89** | **24.23** | **19.39** | **132.09** | **26.49** | **19.27** | **132.79** | **25.65** | **16.46** |
| | *122.57,* | *18.59,* | *25.57,* | *123.42,* | *28.08,* | *16.28,* | *121.41,* | *21.47,* | *16.31,* | *120.02,* | *23.84,* | *16.37,* | *121.59,* | *21.54,* | *13.19,* |
| | *419.54* | *192.01* | *248.80* | *427.50* | *242.72* | *194.66* | *393.61* | *227.49* | *192.95* | *392.42* | *217.32* | *199.01* | *457.32* | *147.41* | *167.66* |
| AB | **105.49** | **21.59** | **21.44** | **106.03** | **21.20** | **19.61** | **107.20** | **28.28** | **19.67** | **103.75** | **19.56** | **16.97** | **103.19** | **10.25** | **19.68** |
| | *83.61,* | *15.82,* | *15.48,* | *84.13,* | *15.47,* | *14.05,* | *84.32,* | *21.53,* | *14.06,* | *82.03,* | *13.91,* | *12.44,* | *82.13,* | *6.01,* | *14.39,* |
| | *259.77* | *106.94* | *117.59* | *260.55* | *97.12* | *116.08* | *245.46* | *134.35* | *115.93* | *251.64* | *103.04* | *107.72* | *262.85* | *39.11* | *108.33* |
| Tu | **153.51** | **35.15** | **44.51** | **153.30** | **41.63** | **36.97** | **152.37** | **30.45** | **36.72** | **157.51** | **48.64** | **36.76** | **143.18** | **32.07** | **27.18** |
| | *118.09,* | *23.38,* | *30.45,* | *117.75,* | *28.48,* | *24.16,* | *116.22,* | *20.82,* | *24.06,* | *120.67,* | *33.70,* | *25.11,* | *110.34,* | *20.19,* | *16.99,* |
| | *307.56* | *79.03* | *98.90* | *305.94* | *96.71* | *81.10* | *301.56* | *75.27* | *80.68* | *312.90* | *106.37* | *82.78* | *287.90* | *75.42* | *64.67* |
| WIK | **152.53** | **20.75** | **45.09** | **154.22** | **57.26** | **11.86** | **151.16** | **28.26** | **12.47** | **144.75** | **38.10** | **14.42** | **157.08** | **70.09** | **16.41** |
| | *124.33,* | *15.31,* | *38.65,* | *126.51,* | *48.69,* | *7.42,* | *123.23,* | *22.63,* | *8.03,* | *116.59,* | *31.88,* | *10.18,* | *130.76,* | *58.90,* | *11.23,* |
| | *438.17* | *152.26* | *353.75* | *448.66* | *415.32* | *107.02* | *425.82* | *220.90* | *113.75* | *381.07* | *291.04* | *132.73* | *513.79* | *463.73* | *145.04* |

**Table S3. Results from analysis of heterogeneity among strains in morphology: performance relationships.** The F-ratio and associated *P*-value (df = 2,81 in all analyses) are presented for the test of the null hypothesis that the strains share the same common slope for that morphological variable on that eigenvector of performance. The morphological traits (rows) are defined in Figure 1; the performance vectors are defined in Table 3. Slopes for this full model are not presented but differences among strains in slope are captured by the results for the subset of best-fit models within each strain (Tables S4-S5).

| | $e_1$ | | $e_2$ | | $e_3$ | |
|---|---|---|---|---|---|---|
| | F-ratio | P-value | F-ratio | P-value | F-ratio | P-value |
| SL | 2.59 | 0.0808$^+$ | 0.27 | 0.7613 | 0.77 | 0.4673 |
| ILD1.12 | 3.13 | 0.0489* | 1.72 | 0.1853 | 0.15 | 0.8600 |
| ILD2.11 | 1.55 | 0.2179 | 0.80 | 0.4528 | 0.47 | 0.6268 |
| ILD2.3 | 0.81 | 0.4473 | 2.58 | 0.0816$^+$ | 0.85 | 0.4317 |
| ILD3.4 | 0.98 | 0.3795 | 1.67 | 0.1939 | 3.17 | 0.0473* |
| ILD3.10 | 0.20 | 0.8206 | 0.75 | 0.4771 | 0.29 | 0.7463 |
| ILD4.5 | 1.44 | 0.2424 | 1.05 | 0.3561 | 1.92 | 0.1534 |
| ILD5.7 | 4.72 | 0.0115* | 2.98 | 0.0562$^+$ | 0.26 | 0.7748 |
| ILD7.8 | 0.49 | 0.6172 | 0.22 | 0.8007 | 0.46 | 0.6328 |
| ILD8.9 | 0.64 | 0.5299 | 0.54 | 0.5857 | 0.52 | 0.5945 |
| ILD9.10 | 3.53 | 0.0337* | 0.53 | 0.5900 | 0.20 | 0.8166 |

* significant at $\alpha = 0.05$
$^+$ significant at $\alpha = 0.1$

**Table S4. Summary of subset of best-fit morphology: performance models per strain for the first major axis of swimming performance, $e_1$.** The number of models (n) in the best-fit subset (i.e., within 2 AIC of the best model), and the minimum and maximum adjusted $R^2$ of that subset is reported in the first column. All traits were included in at least one and up to all models in the subset, as indicated by the % of models in the first row of each section. The average standardised β is reported, along with z-value and *P* for hypothesis test of zero slope, for both averaging across the full subset (i.e., assuming zero slope in models where trait was excluded) and the conditional average over only the sub-subset of models including that predictor. Slopes are bolded where the null hypothesis was rejected (α = 0.05).

| | | SL | ILD1.12 | ILD2.11 | ILD2.3 | ILD3.4 | ILD3.10 | ILD4.5 | ILD5.7 | ILD7.8 | ILD8.9 | ILD9.10 |
|---|---|---|---|---|---|---|---|---|---|---|---|---|
| AB | % models | 4% | 28% | 8% | 32% | 8% | 60% | 100% | 88% | 8% | 44% | 96% |
| | Full subset | | | | | | | | | | | |
| n=25 | β | -0.001 | 0.036 | 0.003 | 0.029 | -0.003 | 0.209 | **0.477** | -0.241 | -0.007 | 0.058 | 0.395 |
| adj R^2: | z-value | 0.049 | 0.339 | 0.079 | 0.274 | 0.074 | 0.694 | 2.063 | 1.165 | 0.137 | 0.47 | 1.894 |
| 15.5 - 22.5% | Pr(>\|z\|) | 0.9613 | 0.7347 | 0.9367 | 0.7838 | 0.9412 | 0.4876 | 0.0391 | 0.2439 | 0.8907 | 0.6381 | 0.0583 |
| | Conditional | | | | | | | | | | | |
| | β | -0.059 | 0.196 | 0.085 | 0.183 | -0.078 | 0.450 | **0.477** | -0.310 | -0.109 | 0.186 | **0.421** |
| | z-value | 0.33 | 1.146 | 0.441 | 0.877 | 0.406 | 1.525 | 2.063 | 1.684 | 0.647 | 1.169 | 2.221 |
| | Pr(>\|z\|) | 0.7413 | 0.2516 | 0.6591 | 0.3806 | 0.6847 | 0.1272 | 0.0391 | 0.0921 | 0.5177 | 0.2424 | 0.0263 |
| Tu | % models | 8% | 100% | 92% | 17% | 17% | 8% | 8% | 100% | 33% | 17% | 100% |
| | Full subset | | | | | | | | | | | |
| n=12 | β | 0.004 | **-0.569** | -0.265 | -0.014 | -0.007 | -0.008 | 0.015 | **0.539** | 0.041 | 0.003 | **0.301** |
| adj R^2: | z-value | 0.09 | 3.411 | 1.401 | 0.235 | 0.145 | 0.134 | 0.208 | 3.609 | 0.375 | 0.045 | 2.382 |
| 49.1 - 53.3% | Pr(>\|z\|) | 0.9283 | 0.0006 | 0.1613 | 0.8146 | 0.8851 | 0.8938 | 0.8349 | 0.0003 | 0.7073 | 0.9640 | 0.0172 |
| | Conditional | | | | | | | | | | | |
| | β | 0.075 | **-0.569** | -0.304 | -0.115 | -0.077 | -0.119 | 0.168 | **0.539** | 0.184 | 0.032 | **0.301** |
| | z-value | 0.406 | 3.411 | 1.771 | 0.853 | 0.531 | 0.610 | 0.940 | 3.609 | 1.100 | 0.146 | 2.382 |
| | Pr(>\|z\|) | 0.6850 | 0.0006 | *0.0765* | 0.3934 | 0.5955 | 0.5422 | 0.3474 | 0.0003 | 0.2713 | 0.8836 | 0.0172 |
| WIK | % models | 100% | 29% | 21% | 7% | 7% | 7% | 79% | 7% | 7% | 7% | 7% |
| | Full subset | | | | | | | | | | | |
| n=14 | β | **0.539** | -0.063 | 0.019 | 0.002 | 0.004 | 0.004 | 0.173 | -0.001 | 0.000 | 0.001 | -0.008 |
| adj R^2: | z-value | 3.633 | 0.443 | 0.209 | 0.055 | 0.098 | 0.105 | 1.014 | 0.04 | 0.003 | 0.022 | 0.163 |
| 30.2 - 34.3% | Pr(>\|z\|) | 0.0003 | 0.6581 | 0.8343 | 0.9562 | 0.9218 | 0.9161 | 0.3104 | 0.9682 | 0.9978 | 0.9821 | 0.8706 |
| | Conditional | | | | | | | | | | | |

| | | | | | | | | | | | |
|---|---|---|---|---|---|---|---|---|---|---|---|
| β | **0.539** | 0.245 | -0.230 | -0.113 | 0.071 | 0.078 | 0.140 | 0.048 | -0.027 | 0.015 | -0.002 |
| z-value | 3.633 | 1.59 | 1.221 | 0.779 | 0.462 | 0.498 | 0.681 | 0.254 | 0.184 | 0.103 | 0.012 |
| Pr(>\|z\|) | 0.0003 | 0.1119 | 0.2222 | 0.4361 | 0.6438 | 0.6187 | 0.4956 | 0.7991 | 0.8541 | 0.9180 | 0.9901 |

**Table S5. Summary of subset of best-fit morphology: performance models per strain for the first major axis of swimming performance, $\boldsymbol{e_2}$.** The number of models (n) in the best-fit subset (i.e., within 2 AIC of the best model), and the minimum and maximum adjusted $R^2$ of that subset is reported in the first column. All traits were included in at least one and up to all models in the subset, as indicated by the % of models in the first row of each section. The average standardised β is reported, along with z-value and *P* for hypothesis test of zero slope, for both averaging across the full subset (i.e., assuming zero slope in models where trait was excluded) and the conditional average over only the sub-subset of models including that predictor. Slopes are bolded where the null hypothesis was rejected (α = 0.05).

| | | SL | ILD1.12 | ILD2.11 | ILD2.3 | ILD3.4 | ILD3.10 | ILD4.5 | ILD5.7 | ILD7.8 | ILD8.9 | ILD9.10 |
|---|---|---|---|---|---|---|---|---|---|---|---|---|
| AB | % models | 0% | 21% | 39% | 3% | 97% | 39% | 3% | 85% | 45% | 30% | 30% |
| | Full subset | | | | | | | | | | | |
| n=33 | β | | 0.039 | 0.121 | 0.001 | -0.307 | -0.065 | -0.001 | 0.252 | -0.089 | 0.057 | -0.040 |
| adj R^2: | z-value | | 0.358 | 0.641 | 0.051 | 1.780 | 0.440 | 0.047 | 1.211 | 0.566 | 0.452 | 0.369 |
| 14.0 - 20.3% | Pr(>\|z\|) | | 0.7207 | 0.5215 | 0.9593 | 0.0751 | 0.6599 | 0.9624 | 0.2258 | 0.5712 | 0.6513 | 0.7122 |
| | Conditional | | | | | | | | | | | |
| | β | | 0.212 | 0.300 | 0.170 | **-0.320** | -0.234 | -0.135 | 0.323 | -0.242 | 0.213 | -0.186 |
| | z-value | | 1.254 | 1.613 | 0.830 | 1.962 | 1.178 | 0.507 | 1.797 | 1.397 | 1.313 | 1.127 |
| | Pr(>\|z\|) | | 0.2098 | 0.1068 | 0.4066 | 0.0498 | 0.2388 | 0.6125 | 0.0723 | 0.1623 | 0.1891 | 0.2596 |
| Tu | % models | 27% | 62% | 19% | 96% | 15% | 4% | 31% | 27% | 15% | 73% | 100% |
| | Full subset | | | | | | | | | | | |
| n=26 | β | -0.057 | -0.133 | 0.041 | -0.335 | -0.009 | 0.000 | -0.051 | -0.056 | 0.000 | -0.191 | **0.382** |
| adj R^2: | z-value | 0.387 | 0.666 | 0.289 | 1.765 | 0.131 | 0.002 | 0.266 | 0.405 | 0.005 | 0.967 | 2.511 |
| 22.7 - 30.6% | Pr(>\|z\|) | 0.6984 | 0.5057 | 0.7728 | 0.0775 | 0.8958 | 0.9981 | 0.7900 | 0.6853 | 0.9961 | 0.3337 | 0.0120 |
| | Conditional | | | | | | | | | | | |
| | β | -0.267 | -0.297 | 0.283 | -0.348 | 0.003 | -0.140 | -0.347 | -0.248 | -0.004 | -0.284 | **0.382** |
| | z-value | 1.250 | 1.483 | 1.061 | 1.917 | 0.013 | 0.592 | 0.910 | 1.291 | 0.015 | 1.602 | 2.511 |
| | Pr(>\|z\|) | 0.2113 | 0.1380 | 0.2885 | 0.0553 | 0.9896 | 0.5537 | 0.3629 | 0.1967 | 0.9878 | 0.1091 | 0.0120 |
| WIK | % models | 10% | 10% | 95% | 48% | 48% | 10% | 52% | 100% | 10% | 10% | 10% |
| | Full subset | | | | | | | | | | | |
| n=21 | β | 0.000 | -0.007 | 0.336 | 0.077 | 0.099 | -0.004 | -0.153 | **-0.411** | 0.004 | -0.006 | 0.001 |
| adj R^2: | z-value | 0.007 | 0.099 | 1.570 | 0.502 | 0.606 | 0.070 | 0.738 | 2.278 | 0.088 | 0.123 | 0.029 |
| 14.6 - 22.1% | Pr(>\|z\|) | 0.9944 | 0.9211 | 0.1164 | 0.6155 | 0.5443 | 0.9439 | 0.4603 | 0.0227 | 0.9297 | 0.9021 | 0.9765 |
| | Conditional | | | | | | | | | | | |

| | | | | | | | | | | | |
|---|---|---|---|---|---|---|---|---|---|---|---|
| β | 0.005 | -0.114 | 0.370 | 0.225 | 0.253 | -0.066 | -0.289 | **-0.411** | 0.066 | -0.093 | 0.021 |
| z-value | 0.031 | 0.446 | 1.901 | 1.188 | 1.482 | 0.316 | 1.417 | 2.278 | 0.394 | 0.557 | 0.131 |
| Pr(>\|z\|) | 0.9752 | 0.6552 | 0.0573 | 0.2350 | 0.1382 | 0.7517 | 0.1564 | 0.0227 | 0.6933 | 0.5778 | 0.8958 |
